# Parvalbumin Interneuron Impairment Leads to Synaptic Transmission Deficits and Seizures in *SCN8A* Epileptic Encephalopathy

**DOI:** 10.1101/2024.02.09.579511

**Authors:** Raquel M. Miralles, Alexis R. Boscia, Shrinidhi Kittur, Shreya R. Vundela, Eric R. Wengert, Manoj K. Patel

## Abstract

*SCN8A* epileptic encephalopathy (EE) is a severe epilepsy syndrome resulting from *de novo* mutations in the voltage-gated sodium channel Na_v_1.6, encoded by the gene *SCN8A*. Na_v_1.6 is expressed in both excitatory and inhibitory neurons, yet previous studies have primarily focused on the impact *SCN8A* mutations have on excitatory neuron function, with limited studies on the importance of inhibitory interneurons to seizure onset and progression. Inhibitory interneurons are critical in balancing network excitability and are known to contribute to the pathophysiology of other epilepsies. Parvalbumin (PV) interneurons are the most prominent inhibitory neuron subtype in the brain, making up about 40% of inhibitory interneurons. Notably, PV interneurons express high levels of Na_v_1.6. To assess the role of PV interneurons within *SCN8A* EE, we used two mouse models harboring patient-derived *SCN8A* gain-of-function mutations, *Scn8a*^D/+^, where the *SCN8A* mutation N1768D is expressed globally, and *Scn8a*^W/+^-PV, where the *SCN8A* mutation R1872W is selectively expressed in PV interneurons. Expression of the R1872W *SCN8A* mutation selectively in PV interneurons led to the development of spontaneous seizures in *Scn8a*^W/+^-PV mice and seizure-induced death, decreasing survival compared to wild-type. Electrophysiology studies showed that PV interneurons in *Scn8a*^D/+^ and *Scn8a*^W/+^-PV mice were susceptible to depolarization block, a state of action potential failure. *Scn8a*^D/+^ and *Scn8a*^W/+^-PV interneurons also exhibited increased persistent sodium current, a hallmark of *SCN8A* gain-of-function mutations that contributes to depolarization block. Evaluation of synaptic connections between PV interneurons and pyramidal cells showed an increase in synaptic transmission failure at high frequencies (80-120Hz) as well as an increase in synaptic latency in *Scn8a*^D/+^ and *Scn8a*^W/+^-PV interneurons. These data indicate a distinct impairment of synaptic transmission in *SCN8A* EE, potentially decreasing overall cortical network inhibition. Together, our novel findings indicate that failure of PV interneuron spiking via depolarization block along with frequency-dependent inhibitory synaptic impairment likely elicits an overall reduction in the inhibitory drive in *SCN8A* EE, leading to unchecked excitation and ultimately resulting in seizures and seizure-induced death.

## Introduction

*SCN8A* epileptic encephalopathy (EE) is a genetic epilepsy syndrome characterized by treatment-resistant seizures, developmental delay, cognitive dysfunction, and an increased incidence of sudden unexpected death in epilepsy (SUDEP).^1–4^ It is caused by *de novo* gain-of-function (GOF) mutations in the *SCN8A* gene,^5^ which encodes the sodium channel Na_v_1.6.^6^ Na_v_1.6 is expressed widely in the central nervous system, and is prominent at the axon initial segment (AIS) of both excitatory and inhibitory neurons.^7–9^ Previous studies using mouse models of *SCN8A* EE show that excitatory neurons are hyperexcitable,^10^ whereas somatostatin inhibitory interneurons experience premature depolarization block, a mechanism of action potential failure.^11^ Despite advances in understanding the physiological mechanisms of *SCN8A* EE, current treatments are often unable to control seizures and reduce the risk of SUDEP, highlighting the need to further understand the underlying network mechanisms of this disorder.

The balance of excitation and inhibition in the brain is critical in seizure generation. Inhibitory interneurons suppress the activity of their target excitatory neurons in an effort to control network dynamics and prevent any excessive excitation that may lead to seizures.^12–15^ Inhibitory interneurons are incredibly diverse; a recent study has identified 28 subtypes based on morphological, electrophysiological, and transcriptomic data.^16^ There are five major subclasses of cortical inhibitory interneurons: parvalbumin (PV), somatostatin (SST), vasoactive intestinal peptide (VIP), Lamp5 (previously 5HT3aR), and Sncg interneurons.^14–17^ The most numerous subtype is PV interneurons, which make up about 40% of inhibitory interneurons and provide feed-forward and feed-back inhibition to networks through reliable, high-frequency firing.^14,15^ PV interneurons are known to express Na_v_1.6 and yet have been previously unstudied in the context of *SCN8A* EE, significantly limiting our understanding of the seizure network in this disorder. Inhibitory interneuron dysfunction has been heavily implicated in Dravet Syndrome, another sodium channelopathy resulting from mutations in the *SCN1A* gene. Previous studies of Dravet Syndrome indicate that PV interneurons are hypoexcitable during a critical developmental time window.^18,19^ In adult mice, PV interneurons show deficits in synaptic transmission and synchronization that likely contribute to the chronic phenotype of Dravet Syndrome.^20,21^ Additionally, PV interneurons have also been implicated in temporal lobe epilepsy (TLE). In mouse models of TLE, previous studies show a reduction in parvalbumin staining, indicating a potential loss of PV interneurons,^22,23^ and others suggest a role for PV interneurons in abnormal synapse formation.^24,25^

In this study, we used two mouse models of *SCN8A* EE harboring the N1768D (*Scn8a*^D/+^) and R1872W (*Scn8a*^W/+^) patient-derived *SCN8A* mutations. These models recapitulate key features of the disease through spontaneous seizures and increased risk of seizure-induced death.^26–28^ *Scn8a*^D/+^ mice express a germline knock-in of the N1768D mutation,^26,27^ whereas *Scn8a*^W/+^ mice harbor a Cre-dependent knock-in of the R1872W mutation.^28^ Previous studies have used this conditional expression model to investigate cell-type specific contributions to *SCN8A* EE: selective expression of the R1872W mutation in forebrain excitatory neurons leads to spontaneous seizures and premature death,^28^ whereas selective expression of this mutation in SST inhibitory interneurons leads to audiogenic seizures without spontaneous seizures or seizure-induced death.^11^

Here, we used both the global *Scn8a*^D/+^ model and the conditional *Scn8a*^W/+^ model of *SCN8A* EE to assess the phenotype of mutant PV interneurons individually and as a component of the *SCN8A* EE network. We report that selective expression of the R1872W *SCN8A* mutation in PV interneurons (*Scn8a*^W/+^-PV) is sufficient to induce spontaneous seizures and premature seizure-induced death, indicating the importance of this inhibitory subtype to *SCN8A* EE as a whole. Whole-cell patch clamp electrophysiology recordings of PV interneurons demonstrated an increased susceptibility to action potential failure via depolarization block. Consequently, we also observed a decrease in spontaneous inhibition received by pyramidal cells in *Scn8a* mutant mice. Recordings of voltage-gated sodium currents showed an elevation of the persistent sodium current (I_NaP_) in both models and an elevation of resurgent sodium current (I_NaR_) in the *Scn8a*^W/+^-PV model, potentially contributing to the premature depolarization block phenotype. Decreased miniature inhibitory postsynaptic currents (mIPSCs) onto *Scn8a*^W/+^-PV pyramidal cells were also observed, suggesting a possible synaptic deficit between PV interneuron and pyramidal cells (PV:PC pairs), and dual recordings of synaptically connected cells revealed an increase in PV:PC synaptic transmission failure at high PV firing frequencies as well as a prolonged synaptic latency. In summary, these data reveal a significant and previously unappreciated impairment of PV interneurons and their synaptic connections to excitatory pyramidal cells in *SCN8A* EE. Selective expression of a *SCN8A* mutation in PV interneurons shows that these impairments are sufficient to cause seizures and SUDEP in mice, indicating the importance of this critical interneuron subtype to seizure generation and redefining our understanding of the cortical microcircuit function in this disease.

## Materials and methods

### Mouse husbandry and genotyping

*Scn8a*^D/+^ and *Scn8a*^W/+^ mice were generated as previously described and maintained through crosses with C57BL/6J mice (Jax, #000664) to keep all experimental mice on a C57BL/6J genetic background.^27,28^ Cell type-specific expression of R1872W was achieved using males heterozygous for the R1872W allele and females homozygous for PV-Cre (Jax, #017320) to generate mutant mice (*Scn8a*^W/+^-PV).^28^ Homozygous PV-IRES-Cre females were used for breeding to ensure minimal germline recombination due to Cre, as shown previously.^29,30^ Because certain transgenic mice entail the insertion of Cre directly into the coding sequence, for all experiments we used WT controls that contained the same Cre allele but lacked the allele encoding the *Scn8a* variant. Fluorescent labeling of PV interneurons was achieved by first crossing male *Scn8a*^D/+^ or *Scn8a*^W/+^ mice with a Cre-dependent tdTomato reporter (Jax, #007909) and then with female mice homozygous for PV-Cre. Experimental groups used ≥3 randomly-selected mice to achieve statistical power and roughly equal numbers of male and female mice to control for potential sex differences. No sex differences were observed. All genotyping was conducted through Transnetyx automated genotyping PCR services.

### *In vivo* seizure monitoring

Custom electroencephalogram (EEG) headsets (PlasticsOne) were implanted in 5-week-old *Scn8a*^W/+^-PV mice using standard surgical techniques as previously described.^31^ Anesthesia was induced with 5% and maintained with 0.5%-3% isoflurane. Adequacy of anesthesia was assessed by lack of toe-pinch reflex. A midline skin incision was made over the skull and connective tissue was removed. Burr holes were made at the lateral/anterior end of the left and right parietal bones to place EEG leads, and at the interparietal bone for ground electrodes. EEG leads were placed bilaterally in the cortex or unilaterally placed in the cortex and superior colliculus using a twist. A headset was attached to the skull with dental acrylic (Jet Acrylic; Lang Dental). Mice received postoperative analgesia with ketoprofen (5 mg/kg, i.p.) and 0.9% saline (0.5 mL i.p.) and were allowed to recover a minimum of 2-5d before seizure-monitoring experiments.

Mice were then individually housed in custom-fabricated chambers and monitored for the duration of the experiment. The headsets were attached to a custom low-torque swivel cable, allowing mice to move freely in the chamber. EEG signals were amplified at 2000× and bandpass filtered between 0.3 and 100 Hz, with an analog amplifier (Neurodata model 12, Grass Instruments). Biosignals were digitized with a Powerlab 16/35 and recorded using LabChart 7 software at 1 kS/s. Video acquisition was performed by multiplexing four miniature night vision-enabled cameras and then digitizing the video feed with a Dazzle Video Capture Device and recording at 30 fps with LabChart 7 software in tandem with biosignals.

### Immunohistochemistry

Brain tissue for immunohistochemistry was processed as previously described.^11,32^ Mice were anesthetized and transcardially perfused with 10 mL Dulbecco’s PBS (DPBS) followed by 10 mL 4% PFA. Brains were immersed in 4% PFA overnight at 4°C and stored in DPBS. 40 μm sagittal brain sections were obtained using a vibratome. Sections were incubated with mouse anti-PV (Millipore) diluted in 2% donkey serum (Jackson ImmunoResearch Laboratories) with 0.1% Triton X (Sigma-Aldrich) at a concentration of 1:1000 in DPBS. The secondary antibody, donkey anti-mouse AlexaFluor-488 (Invitrogen), was diluted 1:1000 in donkey serum (2%) and Triton-X (0.1%) in DPBS. Sections were stained free-floating in primary antibody on a shaker at 4°C overnight and with secondary antibody for 1 h at room temperature the following day. Tissues were counterstained with NucBlue Fixed Cell ReadyProbes Reagent (DAPI) (ThermoFisher Scientific, catalog #R37606) included in the secondary antibody solution. Tissues were mounted on slides using AquaMount (Polysciences).

### Brain Slice Preparation

Preparation of acute brain slices for patch-clamp electrophysiology experiments was modified from standard protocols previously described^10,11,28^. Mice were anesthetized with isoflurane and decapitated. The brains were rapidly removed and kept in chilled ACSF (0°C) containing (in mM): 125 NaCl, 2.5 KCl, 1.25 NaH_2_PO_4_, 2 CaCl_2_, 1 MgCl_2_, 0.5 L-ascorbic acid, 10 glucose, 25 NaHCO_3_, and 2 Na-pyruvate. For dual-cell patch-clamp experiments, the slicing solution was modified to contain (in mM): 93 N-Methyl-D-glucamine (NMDG), 2.5 KCl, 1.25 NaH_2_PO_4_, 20 HEPES, 5 L-ascorbic acid (sodium salt), 2 thiourea, 3 sodium pyruvate, 0.5 CaCl_2_, 10 MgSO_4_, 25 D-glucose, 12 N-acetyl-L-cysteine, 30 NaHCO_3_; pH adjusted to 7.2-7.4 using HCl (osmolarity 310 mOsm). Slices were continuously oxygenated with 95% O_2_ and 5% CO_2_ throughout the preparation. 300 μm coronal or horizontal brain sections were prepared using a Leica Microsystems VT1200 vibratome. Slices were collected and placed in ACSF warmed to 37°C for ∼30 min and then kept at room temperature for up to 6 h.

### Electrophysiology Recordings

Brain slices were placed in a chamber superfused (∼2 ml/min) with continuously oxygenated recording solution warmed to 32 ± 1°C. Cortical layer IV/V PV interneurons were identified as red fluorescent cells, and pyramidal neurons were identified based on morphology and absence of fluorescence via video microscopy using a Carl Zeiss Axioscope microscope. Whole-cell recordings were performed using a Multiclamp 700B amplifier with signals digitized by a Digidata 1322A digitizer. Currents were amplified, lowpass filtered at 2 kHz, and sampled at 100 kHz. Borosilicate electrodes were fabricated using a Brown-Flaming puller (model P1000, Sutter Instruments) to have pipette resistances between 1.5 and 3.5 mΩ. All patch-clamp electrophysiology data were analyzed using custom MATLAB scripts and/or ClampFit 10.7.

### Intrinsic Excitability Recordings

Current-clamp recordings of neuronal excitability were collected in ACSF solution identical to that used for preparation of brain slices. The internal solution contained the following (in mM): 120 K-gluconate, 10 NaCl, 2 MgCl_2_, 0.5 K_2_EGTA, 10 HEPES, 4 Na_2_ATP, 0.3 NaGTP, pH 7.2 (osmolarity 290 mOsm). Intrinsic excitability was assessed using methods adapted from those previously described.^10,11^ Briefly, resting membrane potential was manually recorded from the neuron at rest. Current ramps from 0 to 400 pA over 4 s were used to calculate passive membrane and AP properties, including threshold, upstroke and downstroke velocity, which are the maximum and minimum slopes on the AP, respectively; amplitude, which was defined as the voltage range between AP peak and threshold; APD_50_, which is the duration of the AP at the midpoint between threshold and peak; input resistance, which was calculated using a −20 pA pulse in current-clamp recordings; and rheobase, which was defined as the maximum amount of depolarizing current that could be injected into neurons before eliciting an AP. AP frequency–current relationships were determined using 1s current injections from −140 to 1200 pA. Spikes were only counted if amplitude was >20 mV. The threshold for depolarization block was operationally defined as the current injection step that elicited the maximum number of APs (i.e., subsequent current injection steps of greater magnitude resulted in fewer APs because of entry into depolarization block).

### Outside-Out Voltage-Gated Sodium Channel Recordings

Patch-clamp recordings in the outside-out configuration were collected using a protocol modified from an approach previously described.^10,11^ Recordings were collected in ACSF solution. The internal solution for all voltage-gated sodium channel recordings contained the following (in mM): 140 CsF, 2 MgCl_2_, 1 EGTA, 10 HEPES, 4 Na_2_ATP, and 0.3 NaGTP with the pH adjusted to 7.3 and osmolality to 300 mOsm. Voltage-dependent activation and steady-state inactivation parameters were recorded using voltage protocols previously described.^11^

### Persistent and Resurgent Sodium Current Recordings

The recording solution has been previously described^33,34^ and contained (in mM): 100 NaCl, 40 TEACl, 10 HEPES, 3.5 KCl, 2 CaCl_2_, 2 MgCl_2_, 0.2 CdCl_2_, 4 4-aminopyridine (4-AP), 25 D-glucose. Steady-state persistent sodium currents (I_NaP_) were elicited using a voltage ramp (20 mV/s) from −80 to −20 mV. To record resurgent sodium currents (I_NaR_), PV interneurons were held at −100 mV, depolarized to 30 mV for 20 ms, then stepped to voltages between −100 mV and 0 mV for 40 ms. After collecting recordings at baseline, protocols were repeated in the presence of 1μm tetrodotoxin (TTX; Alomone Labs) to completely isolate I_NaP_ and I_NaR_ currents. TTX-subtracted traces were analyzed by extracting the current at each mV. The half-maximal voltage for activation of I_NaP_ was calculated as previously described.^34^

### Inhibitory Postsynaptic Current Recordings

Patch-clamp recordings of inhibitory postsynaptic currents onto pyramidal cells were performed using the same ACSF external solution and an internal solution containing (in mM): 70 K-Gluconate, 70 KCl, 10 HEPES, 1 EGTA, 2 MgCl_2_, 4 MgATP, and 0.3 Na_3_GTP, with the pH adjusted to 7.2-7.4 and osmolarity to 290 mOsm. Pyramidal cells were held at -70 mV and a 1-min gap-free recording was performed in the current-clamp configuration to assess spontaneous IPSC frequencies before bath application of 500 nM TTX to record miniature IPSCs. After recording spontaneous and miniature IPSCs, 1 μM gabazine was bath applied to block currents and ensure that only inhibitory events were recorded.

### Dual-Cell Synaptic Connection Recordings

Unitary IPSCs (uIPSCs) were obtained via two simultaneous patch-clamp recordings from synaptically-connected neurons located within 50 μm of one another in the somatosensory cortex of a horizontal slice. A 2-ms pulse at 1000 pA elicited action potentials in the presynaptic neuron at 1, 5, 10, 20, 40, 80, and 120 Hz. The internal solution was modified to contain (in mM): 65 K-gluconate, 65 KCl, 2 MgCl_2_, 10 HEPES, 0.5 EGTA, 10 Phosphocreatine-Tris_2_, 4 MgATP, 0.3 NaGTP; pH adjusted to 7.2-7.4 using KOH (osmolarity 290 mOsm).^21^

### Statistical Analysis

Analysis of electrophysiological data was performed blinded. All statistical comparisons were made using the appropriate test in GraphPad Prism 9. Categorical data were analyzed using the Fisher’s exact test. For membrane and AP properties, spontaneous firing frequency, depolarization block threshold, peak sodium currents, half-maximal voltages, IPSC frequency and amplitude, and synaptic uIPSC properties, mouse genotypes were compared by one-way ANOVA followed by Dunnett’s multiple comparisons test when the data were normally distributed with equal variances, by Brown-Forsythe ANOVA with Dunnett’s multiple comparisons test when the data were normally distributed with unequal variances, and by the nonparametric Kruskal–Wallis test followed by Dunn’s multiple comparisons test when the data were not normally distributed. Data were assessed for normality using the Shapiro-Wilk test. Bartlett’s test with p=0.05 was used to assess equal variance. Data were tested for outliers using the ROUT method to identify outliers, and one positive outlier in the *Scn8a*^D/+^ group was not included in analysis of Fig. 5H. No other outliers were identified. A two-way ANOVA followed by Tukey’s test for multiple comparisons was used to compare groups in experiments in which repetitive measures were made from a single cell over various voltage commands or current injections. Cumulative distribution (survival) plots were analyzed by the Log-rank Mantel-Cox test. Data are presented as individual data points and/or mean ± SEM. Exact n and p-values are reported in figure legends.

## Results

### Spontaneous seizures and seizure-induced death in mice with selective expression of mutant Nav1.6 in PV interneurons

We first sought to determine if expression of a gain-of-function *SCN8A* mutation selectively in PV interneurons would be sufficient for the development of spontaneous seizures. We used the conditional knock-in *Scn8a*^W/+^ mouse model and crossed homozygous PV-Cre mice with *Scn8a*^W/+^.tdT mice to generate *Scn8a*^W/+^-PV mice, where the R1872W *SCN8A* mutation is expressed exclusively in PV interneurons (Fig. 1A). *Scn8a*^W/+^-PV mice were implanted with EEG recording electrodes and monitored for 10 weeks. Spontaneous seizures were observed in all recorded *Scn8a*^W/+^-PV mice (*n*=8; Figure 1B). Median seizure onset was about 10 weeks of age. Seizures typically consisted of a wild running phase, which was immediately followed by a tonic-clonic phase in approximately 26% of seizures (23/89). Analysis of EEG signals revealed spike-wave discharges, a distinct aspect of electrographic seizures (Fig. 1C, D). *Scn8a*^W/+^-PV mice also died prematurely compared to WT littermates, with a median survival of 16.6 weeks (Fig. 1E). Electrographic and video recordings confirmed *Scn8a*^W/+^-PV mice that died during monitoring succumbed to seizure-induced death (*n*=3; Supplementary video). Interestingly, all fatal seizures exhibited a tonic phase prior to death, consistent with our previous findings in *SCN8A* EE mice.^35^ Overall, these findings support a previously unappreciated role for PV interneurons in seizure induction and seizure-induced death in a mouse model of *SCN8A* EE.

**Figure 1:**
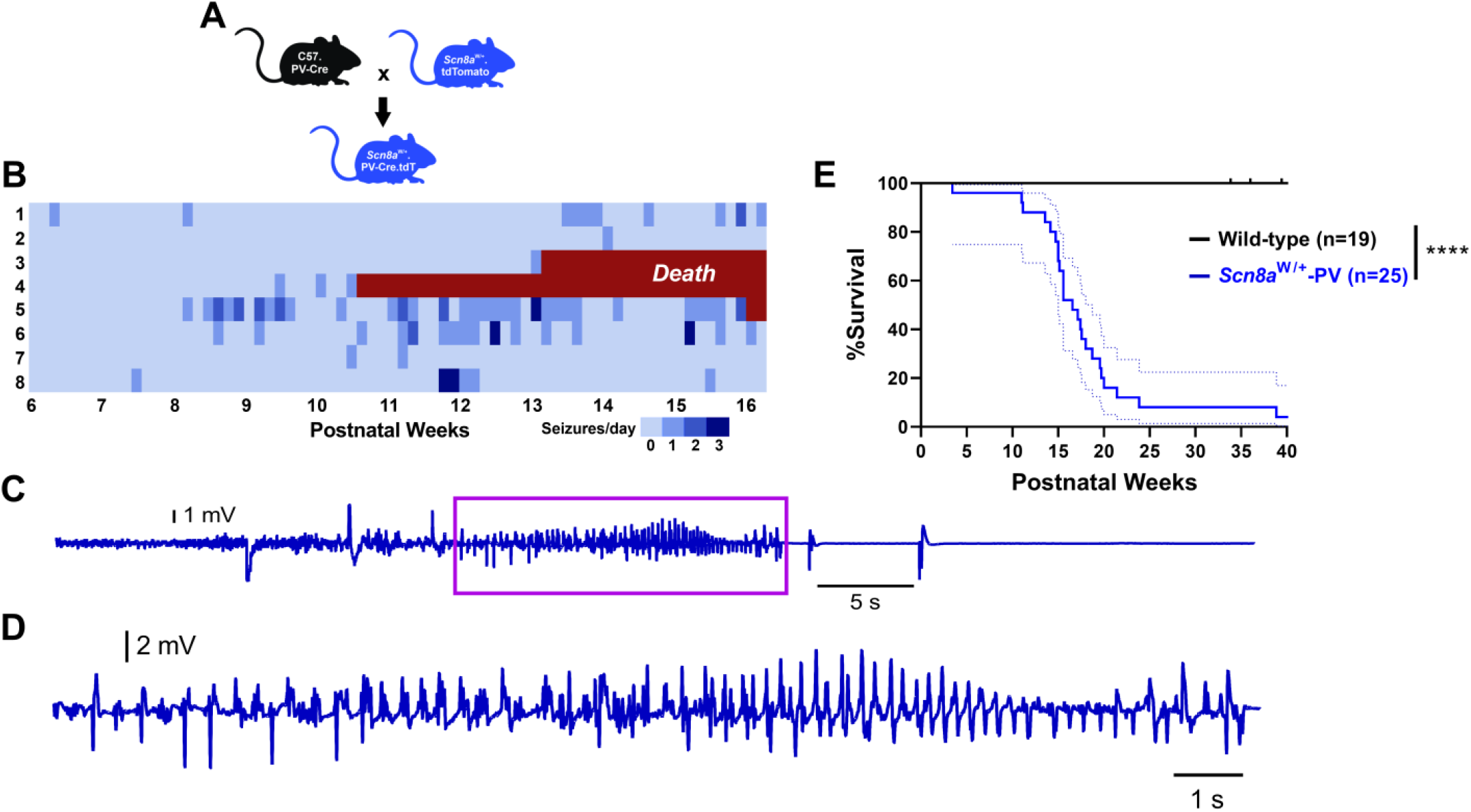
Mice expressing the patient-derived *SCN8A* mutation R1872W exclusively in PV interneurons exhibit spontaneous seizures and seizure-induced death. (**A**) Breeding strategy used to produce *Scn8a*^W/+^.tdT.PV-Cre mice (*Scn8a*^W/+^-PV mice, used for both *in vivo* and whole-cell patch clamp experiments) and age-matched littermate controls on a C57 background. These mice express the R1872W *SCN8A* mutation exclusively in PV interneurons, which are fluorescently labeled with tdTomato. (**B**) Seizure heatmap of (*n*=8) mice over a period of 10 weeks. (**C**) Example spontaneous seizure cortical EEG trace from a *Scn8a*^W/+^-PV mouse. Spontaneous seizure shown here resulted in seizure-induced death (supplementary video). (**D**) Expansion of cortical EEG trace shown in (**C**; purple box) illustrating spike wave discharges. (**E**) Survival of *Scn8a*^W/+^-PV mice (*n*=25) is significantly reduced when compared with WT littermates (*n*=19; *p*<0.0001; Log-rank Mantel-Cox test).

### Depolarization block in *Scn8a* mutant PV interneurons

To assess the intrinsic physiological function of *Scn8a* mutant PV interneurons, we performed electrophysiological recordings of fluorescently labeled PV interneurons in layer IV/V of the somatosensory cortex of adult (5 to 8 weeks) *Scn8a*^D/+^, *Scn8a*^W/+^-PV, and age-matched WT littermates (Fig. 2A, B). WT littermates from both *Scn8a*^D/+^and *Scn8a*^W/+^-PV genotypes did not exhibit any differences in firing frequencies (*p*=0.656) and were pooled. Analysis of membrane and action potential (AP) properties revealed that *Scn8a*^D/+^ PV interneurons had decreased downstroke velocity as well as increased AP width when compared to WT (Table 1). Using a series of depolarizing current injection steps to assess intrinsic excitability, we observed a difference in excitability (*p*=0.028) between WT, *Scn8a*^D/+^, and *Scn8a*^W/+^ PV interneurons. Initially, PV interneurons expressing either *Scn8a* mutation were hyperexcitable compared to WT littermates at lower current injection steps (<100 pA in *Scn8a*^D/+^ mice, *p*=0.045 and <360 pA in *Scn8a*^W/+^-PV mice, *p*=0.030). However, at higher current injection steps, both *Scn8a*^D/+^, and *Scn8a*^W/+^ PV interneurons exhibited progressive action potential failure as a result of depolarization block (>640 pA in *Scn8a*^D/+^ mice, *p*=0.042; >840 pA in *Scn8a*^W/+^-PV mice, *p*=0.041; Fig. 2C-G). Both *Scn8a*^D/+^ and *Scn8a*^W/+^ PV interneurons were more prone to depolarization block than their WT counterparts over the range of current injection magnitudes (*p*<0.0001 and *p*=0.016 respectively; Fig. 2G). Depolarization block of inhibitory interneurons has been previously implicated in seizure-like activity both *in vitro* and *in vivo*, and has been proposed as a biophysical mechanism underlying approach of seizure threshold.^36–41^ Here, the early onset of depolarization block in *Scn8a* mutant PV interneurons indicates a PV hypo-excitability phenotype, similar to the phenotypes observed in PV interneurons in gain-of-function *SCN1A* EE and in SST interneurons in *SCN8A* EE.^11,36^

**Figure 2:**
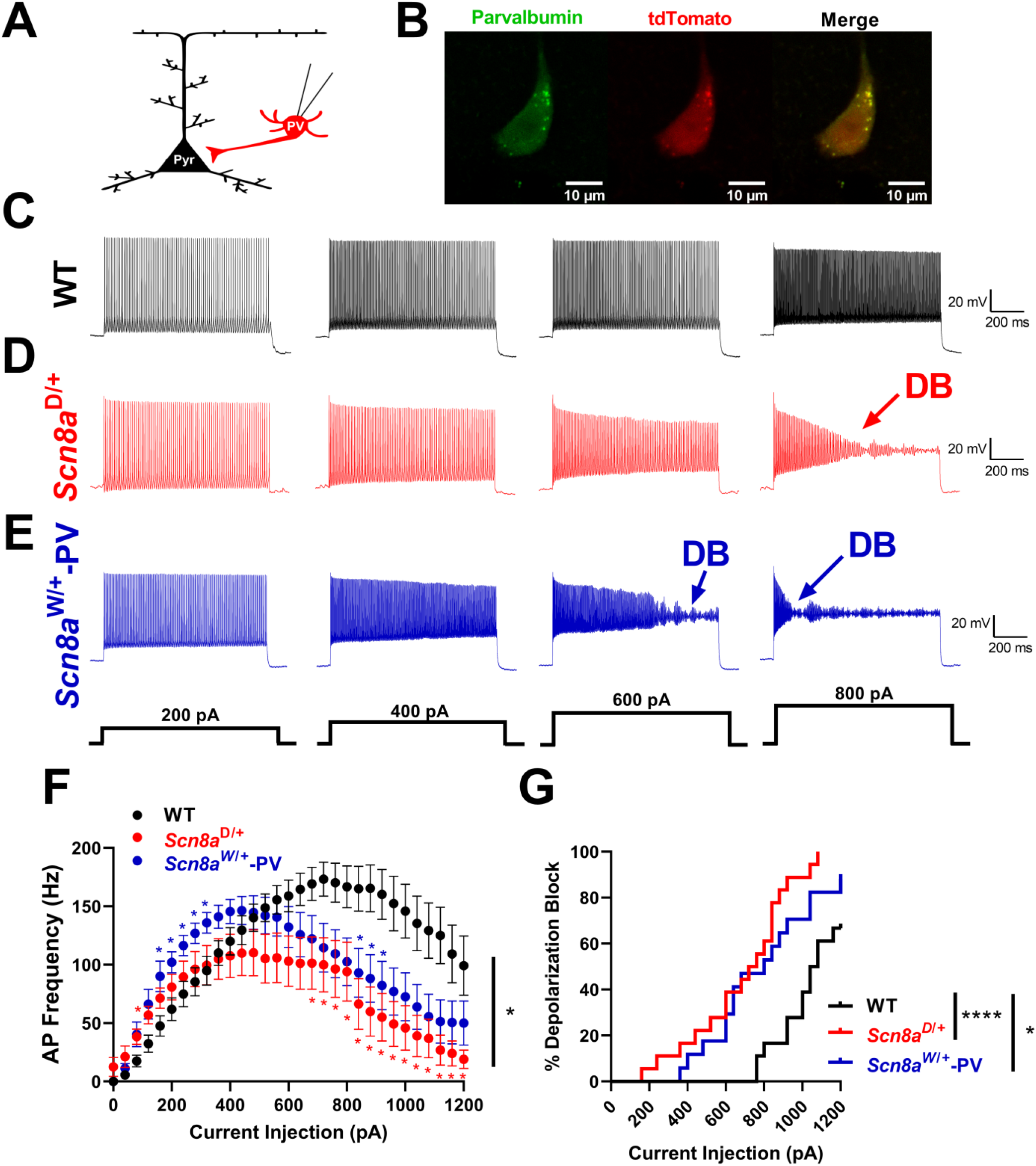
Altered excitability and depolarization block in *Scn8a*^D/+^ and *Scn8a*^W/+^-PV interneurons. (**A**) Whole-cell recordings were collected from WT, *Scn8a*^D/+^, and *Scn8a*^W/+^-PV interneurons in mouse layer IV/V somatosensory cortex. (**B**) Example immunohistochemistry images showing colocalization of parvalbumin (green) and tdTomato (red) immunofluorescence. Scale bar 10μM. (**C-E**) Example traces of WT (**C**, black), *Scn8a*^D/+^ (**D**, red), and *Scn8a*^W/+^ (**E**, blue) PV interneuron firing at 200, 400, 600, and 800 pA current injections. Depolarization block is noted with arrows (DB). (**F**) *Scn8a*^D/+^ (*n*=17, 6 mice) and *Scn8a*^W/+^ PV (*n*=17, 5 mice) interneurons experience a decrease in firing via depolarization block (*, *p*<0.05, two-way ANOVA with Tukey’s multiple comparisons test) when compared to WT PV interneurons (*n*=18, 8 mice). Red or blue stars indicate individual points of significance for either *Scn8a*^D/+^ or *Scn8a*^W/+^-PV, respectively, by multiple comparisons test. (**G**) Cumulative distribution of PV interneuron entry into depolarization block relative to current injection magnitude for WT, *Scn8a*^D/+^, and *Scn8a*^W/+^-PV mice (****, *p*<0.0001, *, *p*<0.05, log-rank Mantel-Cox test).

Previous studies show that pyramidal neurons in global knock-in *Scn8a*^D/+^ mice are hyperexcitable compared to WT, suggesting that a global change in neuronal activity of both inhibitory and excitatory neurons likely contributes to the seizure phenotype.^10^ To determine if firing is affected in excitatory neurons from *Scn8a*^W/+^-PV mice, which selectively express an *Scn8a* mutation in PV interneurons, we recorded the intrinsic excitability of pyramidal neurons from cortical layers IV/V. Interestingly, we did not observe any differences in the intrinsic excitability of pyramidal neurons between the WT and *Scn8a*^W/+^-PV genotypes (*p*=0.446). This suggests that alterations in the physiology of PV interneurons may be sufficient in facilitating seizures in *SCN8A* EE.

### Gain-of-function Nav1.6 mutations impact sodium channel currents in PV interneurons

Depolarization block in *Scn8a*^D/+^ and *Scn8a*^W/+^ PV interneurons likely arises from abnormal sodium channel activity as a result of the gain-of-function mutation, contributing to changes in membrane depolarization levels and subsequent sodium channel availability for AP initiation. Increases in the persistent sodium channel current (I_NaP_) have been identified as a major factor in many epileptic encephalopathy-causing mutations, including both the N1768D and R1872W mutations in *SCN8A* EE^5,10,28,42^ Further, I_NaP_ is a known determinant of depolarization block threshold.^11^ In view of this, we recorded I_NaP_ in PV interneurons. I_NaP_ was increased in both *Scn8a*^D/+^ (−293.1 ± 38.0 pA; *p*=0.032) and *Scn8a*^W/+^ (−347.1 ± 49.0 pA; *p*=0.004) PV interneurons when compared to WT (−166.6 ± 29.7 pA; Fig. 3A-D). Half-maximal voltage of activation (V_1/2_) did not differ from WT (−62.0 ± 1.0 mV) in either *Scn8a*^D/+^ (−59.9 ± 1.1 mV; *p*=0.301) or *Scn8a*^W/+^-PV (- 63.9 ± 1.2 mV; *p*=0.410) mice. Another component of the sodium current that may affect excitability particularly in fast spiking cells is the resurgent sodium current (I_NaR_).^43,44^ I_NaR_ is a slow inactivating depolarizing current that can contribute to increased AP frequency by providing additional depolarization during the falling phase of an AP.^43–45^ I_NaR_ has been previously implicated in temporal lobe epilepsy as well as in sodium channelopathies.^46,47^ I_NaR_ was significantly increased in *Scn8a*^W/+^-PV interneurons (−1136.0 ± 178.5 pA; *p*=0.037), and while we observed an increasing trend, I_NaR_ was not significantly increased in *Scn8a*^D/+^ PV interneurons (−952.8 ± 172.9 pA; *p*=0.219), when compared to WT (−595.9 ± 84.8 pA) PV interneurons (Fig. 3E-H). These results demonstrate an increase in two components of the overall sodium current in PV interneurons, which possibly contributes to their initial hyperexcitability and premature depolarization block. Increases in both I_NaP_ and I_NaR_ likely provide a sustained level of depolarization, resulting in the accumulation of inactivated sodium channels and premature depolarization block. ^11,36,48^

**Figure 3:**
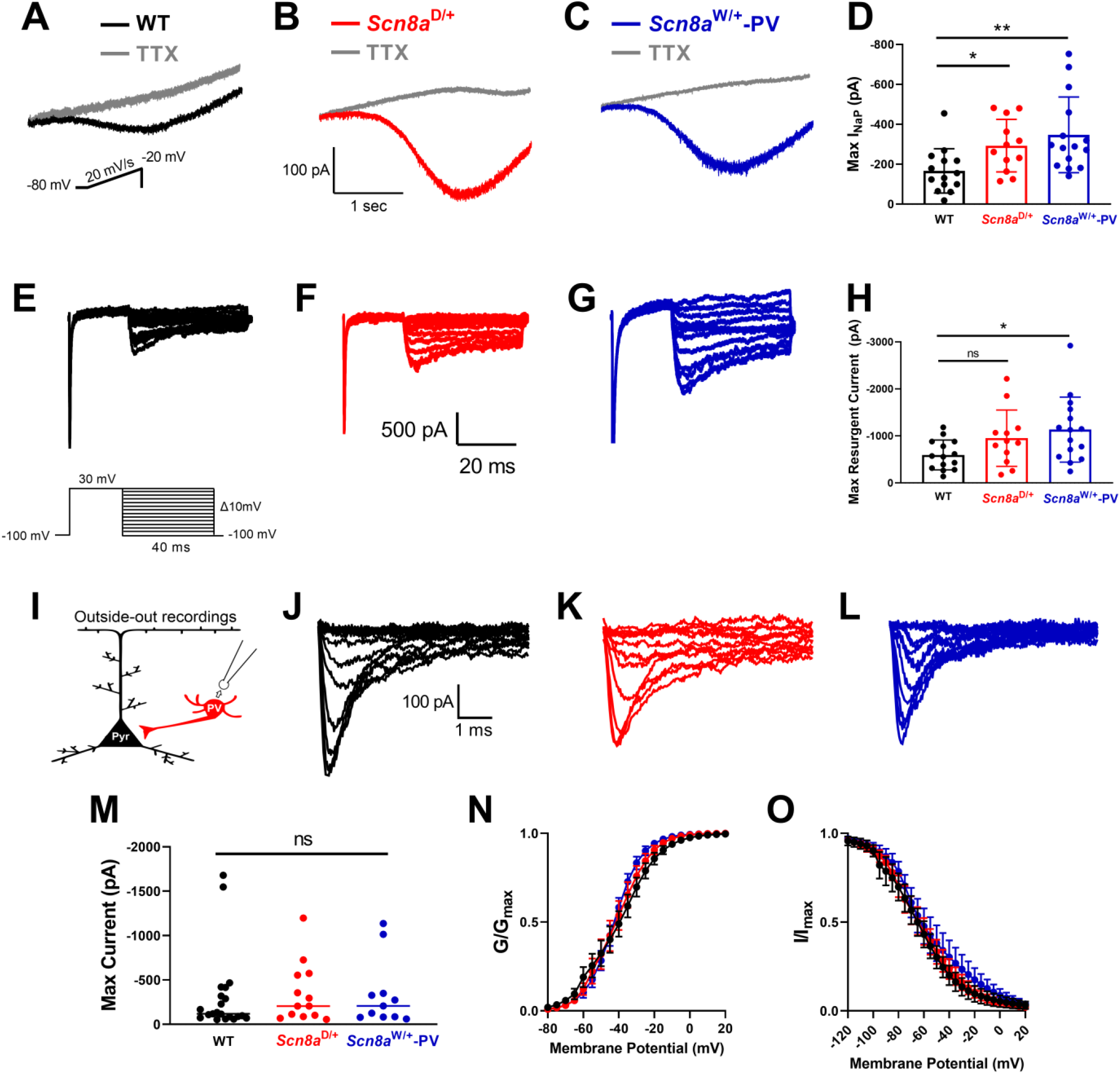
Voltage-gated sodium currents in WT, *Scn8a*^D/+^, and *Scn8a*^W/+^ PV interneurons. (**A, B, C**) Example traces of steady state I_NaP_ evoked by slow voltage ramps from WT (**A**, black), *Scn8a*^D/+^ (**B**, red), and *Scn8a*^W/+^ (**C**, blue) PV interneurons. Traces in gray show slow voltage ramp in the presence of 500 nM TTX. (**D**) Elevated maximum I_NaP_ in *Scn8a*^D/+^ (*n*=12, 4 mice, *, *p*<0.05) and *Scn8a*^W/+^ (*n*=15, 5 mice, **, *p*<0.01) PV interneurons compared with WT (*n*=14, 5 mice) PV interneurons (Kruskal-Wallis test with Dunn’s multiple comparison test). (**E-G**) Example traces of TTX-subtracted I_NaR_ for WT (**E**, black), *Scn8a*^D/+^ (**F**, red), and *Scn8a*^W/+^ (**G**, blue) evoked by voltage commands in which the cell was stepped to membrane potentials of -100 to 0mV, increments of 10mV for 40 ms after first being stepped to 30mV for 20ms. (**H**) Maximum I_NaR_ magnitude was increased between WT (*n*=14, 5 mice) and *Scn8a*^W/+^-PV (*n*=15, 5 mice) interneurons (*, *p*<0.05, Brown-Forsythe ANOVA with Dunnett’s multiple comparison test), whereas I_NaR_ magnitude between WT and *Scn8a*^D/+^ (*n*=12, 4 mice) PV interneurons was not significantly different (*p*<0.05, Brown-Forsythe ANOVA with Dunnett’s multiple comparison test). (**I**) Somatic sodium current was assessed in PV interneurons in-slice using patch-clamp recordings in the outside out configuration. (**J-L**) Example traces for family of voltage-dependent sodium currents recorded from outside-out excised patches from WT (**J**, black), *Scn8a*^D/+^ (**K**, red), and *Scn8a*^W/+^ (**L**, blue) PV interneurons. (**M**) Maximum transient sodium current was not significantly different between WT (*n*=20, 8 mice), *Scn8a*^D/+^ (*n*=12, 4 mice), and *Scn8a*^W/+^ (*n*=12, 4 mice) PV interneurons (*p*>0.05, Kruskal-Wallis test with Dunn’s multiple comparison test). (**N**) Voltage-dependent conductance curve does not differ significantly between WT, *Scn8a*^D/+^, and *Scn8a*^W/+^ PV interneurons (*p*>0.05, 2-way ANOVA). (**O**) Steady-state inactivation does not differ significantly between WT (*n*=12, 4 mice), *Scn8a*^D/+^ (*n*=11, 4 mice), and *Scn8a*^W/+^ (*n*=10, 4 mice) PV interneurons (*p*>0.05, 2-way ANOVA). Boltzmann curves shown are the average of individual curves generated from fits to data points.

Alterations of both activation and steady-state inactivation parameters of the transient sodium channel current have been previously reported in cells expressing GOF *SCN8A* mutations.^5,49–51^ To examine PV interneuron sodium channel currents, we performed excised somatic patches in the outside-out configuration from PV interneurons (Fig. 3I). Sodium current density, voltage-dependent activation, or steady-state inactivation were not different between WT, *Scn8a*^D/+^, and *Scn8a*^W/+^-PV mice (Fig. 3 J-O; Table 2).

### Decreased inhibitory input onto excitatory neurons in *Scn8a* mutant mice

Impaired excitability in *Scn8a* mutant PV interneurons may lead to decreased inhibition onto excitatory pyramidal cells, as PV interneurons are known to directly inhibit pyramidal cells at the soma or AIS.^14,15^ To examine how alterations in PV interneuron excitability affect the cortical network, we recorded spontaneous and miniature inhibitory post-synaptic currents (sIPSCs and mIPSCs) from pyramidal cells (Fig. 4A) as a functional indicator of PV interneuron activity and connectivity. We found that pyramidal cells receive significantly fewer sIPSCs in both *Scn8a*^D/+^ (4.22 ± 0.64 Hz; *p*=0.029) and *Scn8a*^W/+^-PV (3.68 ± 0.95 Hz; *p*=0.002) mice than their WT counterparts (7.97 ± 0.88 Hz; Fig. 4B-C), suggesting a decrease in inhibitory input onto pyramidal cells. sIPSC frequencies between *Scn8a*^D/+^ and *Scn8a*^W/+^-PV pyramidal cells were not different (*p*>0.99), which may imply that PV interneurons are largely responsible for the decrease in somatic inhibitory input in the global *Scn8a*^D/+^ model. sIPSC amplitude was not different between WT (−62.7 ± 4.3 pA), *Scn8a*^D/+^ (−64.4 ± 9.1 pA), and *Scn8a*^W/+^-PV mice (−57.6 ± 9.2 pA; *p*=0.270, Fig. 4D). sIPSC recordings include both AP-induced synaptic transients as well as mIPSCs, which occur due to spontaneous vesicle fusion in the absence of an AP.^52,53^ To isolate AP-independent events, we performed recordings in the presence of TTX (500 nM). Relative to WT controls (3.64 ± 0.65 Hz), we found no significant difference in pyramidal cell mIPSC frequency in *Scn8a*^D/+^ mice (2.79 ± 0.58 Hz; *p*=0.416), but we did observe a significant reduction of mIPSC frequency in *Scn8a*^W/+^-PV mice (1.59 ± 0.26 Hz; *p*=0.017; Fig. 4E), which could underlie impaired synaptic transmission in *Scn8a*^W/+^-PV mice. mIPSC amplitude did not differ between WT (−40.8 ± 3.5 pA), *Scn8a*^D/+^ (−44.1 ± 3.3 pA), and *Scn8a*^W/+^-PV mice (−50.2 ± 9.7 pA; *p*=0.578, Fig. 4F).

**Figure 4:**
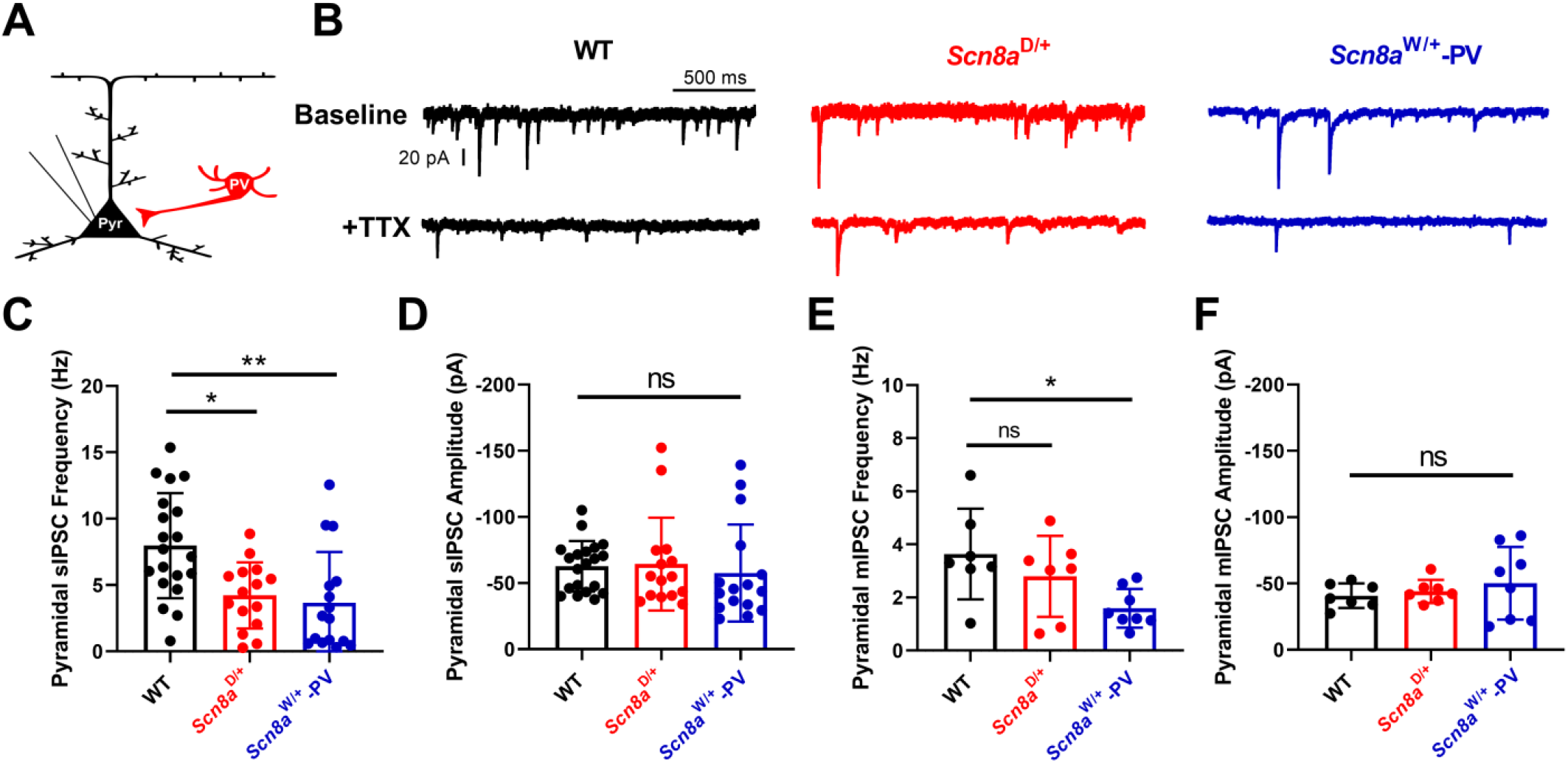
Inhibitory post-synaptic currents (IPSCs) onto pyramidal cells in WT, *Scn8a*^D/+^, and *Scn8a*^W/+^-PV mice. (**A**) Whole-cell recordings of IPSCs were collected from cortical layer V pyramidal cells in WT, *Scn8a*^D/+^, and *Scn8a*^W/+^-PV mice. (**B**) Example traces of IPSCs onto pyramidal cells from WT (black), *Scn8a*^D/+^ (red), and *Scn8a*^W/+^-PV (blue) mice. (**C**) Frequency of sIPSCs onto pyramidal cells is decreased in *Scn8a*^D/+^ (*n*=15, 4 mice, *, *p*<0.05) and *Scn8a*^W/+^-PV (*n*=16, 5 mice, **, *p*<0.01) mice when compared to WT (*n*=20, 6 mice, Kruskal-Wallis test with Dunn’s multiple comparison test). (**D**) Amplitude of sIPSCs onto pyramidal cells is not significantly different between groups (*p*>0.05, Kruskal-Wallis test). (**E**) Frequency of mIPSCs onto pyramidal cells is decreased in *Scn8a*^W/+^-PV (*n*=8, 3 mice) mice when compared to WT (*n*=7, 3 mice, *, *p*<0.05), whereas frequency of mIPSCs onto pyramidal cells in *Scn8a*^D/+^ (*n*=7, 3 mice) mice did not significantly differ from WT (*p*>0.05, one-way ANOVA with Dunnett’s multiple comparison test). (**F**) Amplitude of mIPSCs onto pyramidal cells is not significantly different between WT, *Scn8a*^D/+^, and *Scn8a*^W/+^-PV mice (*p*>0.05, Brown-Forsythe ANOVA with Dunnett’s multiple comparison test).

**Figure 5:**
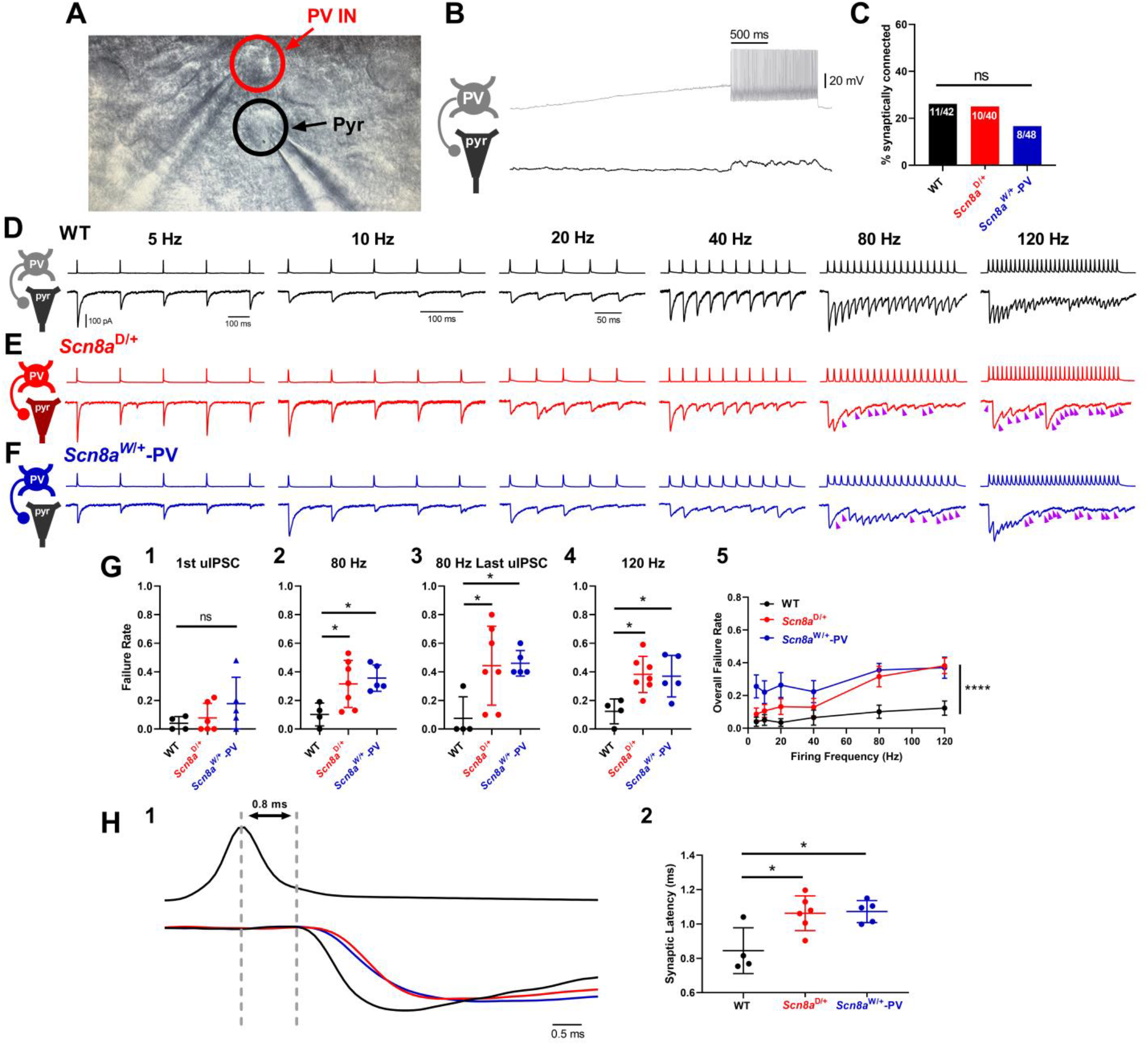
Increased synaptic transmission failure and synaptic latency in *Scn8a* mutant mice. (**A**) Image of dual whole-cell recording of a synaptically connected PV interneuron and pyramidal cell pair. (**B**) Example traces from a PV interneuron (grey) and synaptically coupled pyramidal cell (black). (**C**) Proportion of successfully patched PV:PC pairs that were synaptically connected did not differ between WT (42 pairs from 10 mice), *Scn8a*^D/+^ (40 pairs from 8 mice), and *Scn8a*^W/+^-PV (48 pairs from 7 mice). (**D-F**) Example of presynaptic firing and evoked uIPSCs in WT (**D**), *Scn8a*^D/+^ (**E**), and *Scn8a*^W/+^-PV (**F**) connected pairs at 5 Hz, 10Hz, 20 Hz, 40 Hz, 80 Hz, and 120 Hz. Purple arrows denote uIPSC failures in the postsynaptic neuron. (**G**) Summary data for failure rates of evoked uIPSCs at various frequencies. uIPSC failure rate is not significantly different for the first evoked uIPSC (**G1**, *p*>0.05, Kruskal-Wallis test with Dunn’s multiple comparison test) but is significantly higher in *Scn8a*^D/+^ (*n*=7, 5 mice) and *Scn8a*^W/+^-PV (*n*=5, 4 mice) pairs than WT (*n*=4, 3 mice) at PV interneuron firing frequencies of 80 and 120 Hz (**G2, G4**, *, *p*<0.05, one-way ANOVA with Dunnett’s multiple comparison test). Failure rate of the last uIPSC is higher at 80Hz stimulation frequency (**G3**, *, *p*<0.05, Kruskal-Wallis test with Dunn’s multiple comparison test). Overall failure rate is higher in *Scn8a*^D/+^ and *Scn8a*^W/+^-PV mice (**G5**, ****, *p*<0.0001, two-way ANOVA with Tukey’s multiple comparison test). **(H)** Example traces illustrating synaptic latency in WT, *Scn8a*^D/+^, and *Scn8a*^W/+^-PV, measured from the peak of the presynaptic AP to the onset of the evoked uIPSC (**H1**). Grey dotted lines indicate this latency in WT. Latency is increased in *Scn8a*^D/+^ and *Scn8a*^W/+^-PV mice (**H2**, one-way ANOVA with Dunnett’s multiple comparison test, *, *p*<0.05).

### PV interneuron synaptic transmission is impaired in *Scn8a* mutant mice

Impairment of synaptic transmission has been suggested as a disease mechanism in multiple epilepsy syndromes, notably Dravet Syndrome,^21,54,55^ and proper synaptic signaling is tightly linked to sodium channel function.^56^ To assess how Na_v_1.6 function influences PV interneuron-mediated inhibitory synaptic transmission, we performed dual whole-cell patch clamp recordings of PV interneurons and nearby pyramidal cells (PCs) to find synaptically-connected pairs of cells (Fig. 5A). Synaptically-connected pairs were identified using a current ramp in the presynaptic PV interneuron to elicit inhibitory postsynaptic potentials (IPSPs) in the postsynaptic PC corresponding to each AP in the PV interneuron (Fig. 5B). The number of synaptically-connected PV:PC pairs relative to the total number of pairs was not significantly different between WT, *Scn8a*^D/+^, and *Scn8a*^W/+^-PV mice (*p*=0.309, Fig. 5C). In PV:PC connected pairs, we measured the properties of unitary inhibitory postsynaptic currents (uIPSCs) in PCs evoked by stimulation of PV interneurons. To accurately detect uIPSCs, a high chloride internal solution was used to allow recording of uIPSCs as large inward currents and IPSPs as large membrane depolarizations, overall minimizing the possibility of inaccurately reporting a synaptic failure.

Previous studies indicate that the PV:PC synapse is extremely reliable since PV interneurons have multiple synaptic boutons and a high release probability, indicative of a highly stable synapse.^57^ PV interneurons are also known to fire reliably at high frequencies.^15^ We found that stimulation of PV interneurons at a 1Hz frequency reliably initiated single action potentials in WT mice. Although we detected some failures in *Scn8a*^D/+^ and *Scn8a*^W/+^-PV mice, there was no significant difference in synaptic failure at a frequency of 1Hz (*p*=0.317; Fig. 5G-1) between the groups, suggesting no deficit in synaptic transmission at low stimulation frequencies. The amplitudes of the uIPSCs also did not differ between genotypes (Table 3, *p*=0.427). Additionally, to identify any deficits in short-term synaptic plasticity, we used the first two IPSCs (IPSC1 and IPSC2) elicited by a presynaptic action potential to quantify the paired-pulse ratio (PPR). The PV:PC synapse is known to experience short-term plasticity through synaptic depression.^58,59^ We observed synaptic depression in WT, *Scn8a*^D/+^, and *Scn8a*^W/+^-PV connected pairs, with no significant difference in PPR between WT and *Scn8a* mutant pairs (Table 3).

To analyze activity-dependent synaptic failure, we then used stimulation trains to elicit multiple action potentials at increasing frequencies (5, 10, 20, 40, 80, and 120 Hz; Fig. 5D-F, Table 3). At each frequency, we measured the failure rate of the first and last uIPSC as well as the overall failure rate. Failure rate of the first uIPSC remained low and consistent between WT, *Scn8a*^D/+^, and *Scn8a*^W/+^-PV mice. At lower frequencies (≤40 Hz), there were no differences in overall failure rate or last uIPSC failure rate between WT, *Scn8a*^D/+^, and *Scn8a*^W/+^-PV mice, however, failure rates were increased at frequencies ≥80 Hz. At 80Hz, the overall failure rate in a 20-pulse train was increased in both *Scn8a*^D/+^ (0.316 ± 0.062; *p*=0.035) and *Scn8a*^W/+^-PV (0.356 ± 0.041; *p*=0.020) mice compared to WT (0.100 ± 0.040; Fig. 5G-2), with failures occurring approximately three times as frequently in *Scn8a*^W/+^ mutant mice when compared to WT. Similarly, at 120 Hz stimulation frequency with a 30-pulse train, failure rates observed in *Scn8a*^D/+^ and *Scn8a*^W/+^-PV pairs were greater (0.347 ± 0.095 and 0.369 ± 0.145 respectively; *p*=0.010 and *p*=0.021) than those observed in their WT counterparts (0.123 ± 0.087; Fig. 5G-4). The progression of total activity-dependent synaptic failure through increasing presynaptic stimulation frequencies is shown in Fig. 5G. Additionally, synaptic failure of the last uIPSC in a stimulation train occurred in >40% of trials on average with a stimulation frequency of 80 or 120 Hz. We observed that this increase in synaptic failure is significant for the last uIPSC in an 80 Hz train in *Scn8a*^D/+^ (*p*=0.027) and *Scn8a*^W/+^-PV (*p*=0.033; Fig. 5G-3), supporting a greater degree of activity-dependent failure. Analysis of synaptic latency times, measured from the peak of the presynaptic action potential to the onset of the postsynaptic uIPSC, revealed an increase in synaptic latency in *Scn8a*^D/+^ (*p*=0.010) and *Scn8a*^W/+^-PV (*p*=0.010) mice when compared to WT mice (Fig. 5H; Table 3). Prolonged synaptic latency would suggest an impairment in conduction velocity or GABA release probability, potentially with a longer time lag to vesicle release.^60–63^ Efficient synaptic transmission and vesicle release is critical for overall network inhibition.^64^

## Discussion

Parvalbumin interneurons prominently express Na_v_1.6^8,9^ and are known to play a major role in various epilepsies.^18,21–25,65^ However, their role in the pathophysiology of *SCN8A* epileptic encephalopathy is unknown. Here, we show that: (1) expression of the patient-derived R1872W *SCN8A* GOF mutation selectively in PV interneurons conveys susceptibility to spontaneous seizures and premature seizure-induced death; (2) GOF *SCN8A* mutations in PV interneurons lead to initial hyperexcitability and subsequent action potential failure via depolarization block; (3) PV interneurons in both GOF *SCN8A* mouse models exhibit epileptiform increases in I_NaP_ currents that would facilitate depolarization block; (4) inhibitory input onto excitatory pyramidal cells is significantly reduced in *Scn8a* mutant mice; and (5) there is a progressive, activity-dependent increase in synaptic transmission failure from PV inhibitory interneurons onto excitatory neurons.

Our findings highlight a newfound role for PV interneurons in the pathophysiology of seizures and seizure-induced death in a mouse model of *SCN8A* epileptic encephalopathy.

### Expression of R1872W *SCN8A* mutation in PV interneurons is sufficient to cause seizures and premature death

PV interneurons are known to be the main drivers for seizure activity in Dravet Syndrome, a disorder characterized by deficits in inhibitory neurons, primarily due to haploinsufficiency of Na_v_1.1.^18–21^ Selective deletion of Na_v_1.1 in PV interneurons leads to reduced PV interneuron excitability, decreased spontaneous inhibition of excitatory neurons, and increased susceptibility to seizures.^65^ Similar to impairments observed in mouse models of Dravet Syndrome, we show here that selective expression of the GOF R1872W mutation in PV interneurons is sufficient to induce spontaneous seizures and leads to seizure-induced death (SUDEP) in mice. These findings not only support an important role for PV interneurons in the seizure phenotype of *SCN8A* EE but also provide support for a major role for Na_v_1.6 channels in controlling PV interneuron excitability in addition to Na_v_1.1 channels.

### Gain-of-function *SCN8A* mutations result in premature PV interneuron depolarization block

Proper function of *Scn8a* is critical in repetitive firing,^45^ and as such, we reasoned that mutations affecting the function of *Scn8a* would impact the high-frequency, repetitive firing characteristic of PV interneurons. However, although at lower current injection magnitudes PV interneurons from both mutant mouse models were hyperexcitable, at higher magnitudes we observed PV interneuron action potential failure through depolarization block, resulting in overall PV interneuron hypoexcitability. Depolarization block due to a GOF sodium channel mutation has been shown previously in both *SCN8A* EE and *SCN1A* epileptic encephalopathy^11,36^ Additionally, depolarization block in PV interneurons leads to hyperactivity and subsequent epileptic discharges in excitatory cells, and rescue of depolarization block via optogenetic stimulation leads to a reduction in epileptiform activity.^38–40^ Further, *in vivo* recording of PV interneurons shows evidence for PV depolarization block during seizure activity.^41^ Importantly, this depolarization block and subsequent hypoexcitability in inhibitory interneurons indicates that a mechanism for seizures in *SCN8A* EE, a disorder characterized by primarily GOF sodium channel mutations, may show significant similarities to that of Dravet Syndrome, a disorder primarily characterized by sodium channel haploinsufficiency in inhibitory neurons.

### Impaired synaptic transmission between mutant PV interneurons and pyramidal cells

Our study is the first to examine alterations in synaptic transmission between PV interneurons and excitatory neurons in *SCN8A* EE, and we show a distinct impairment of inhibitory synaptic transmission onto excitatory pyramidal cells in two patient-derived mutation models. Synaptic transmission in *Scn8a* mutant mice failed in an activity-dependent manner, which, considering the fast-spiking nature of PV interneurons and the degree of inhibitory input they provide on neuronal excitatory networks, could have a significant impact on overall seizure susceptibility. A likely mechanism for this failure could be impaired AP propagation, as proper signaling from PV interneurons requires a specific density and function of sodium channels.^56^ This is further supported by the observed increase in synaptic latency, indicating that propagation may be slowed in *SCN8A* mutant mice. Similarly, synaptic transmission between PV interneurons and pyramidal cells is also impaired in Dravet Syndrome, although unlike our findings in *SCN8A* EE, intrinsic excitability deficits are restored in adult PV interneurons.^19,21^

Both depolarization block and synaptic transmission failure occurred at high PV interneuron firing frequencies, and as such, it is important to consider *in vivo* firing frequencies of PV interneurons. PV interneurons are a heterogeneous group made up of primarily basket cells and chandelier cells, which are named for their unique morphologies. These subtypes have slightly different firing patterns and synaptic targets.^14,15^ Since our recordings are focused within cortical layers IV/V, it is likely that we recorded primarily from PV-positive basket cells rather than chandelier cells. *In vivo*, PV interneurons, particularly basket cells, are phase-locked to gamma oscillations, which typically occur between 40-100 Hz.^66^ Events such as sharp wave ripples (SWRs) can lead to PV firing frequencies of >120Hz *in vivo*.^67^ This demonstrates the relevance of both premature PV interneuron depolarization block and activity-dependent PV:PC synaptic transmission failure with expression of mutant *Scn8a*. These gamma oscillations and SWRs are most often associated with the hippocampus, however, there is evidence for oscillations in the cortex.^68,69^ Recently, SWRs have been associated with epileptic discharges in Dravet Syndrome: an increase in SWR amplitude may lead to inhibitory depolarization block and a shift into seizure-like activity.^70^ Considering the frequencies at which we observe PV interneuron failure in *Scn8a* mutant mice, depolarization block and activity-dependent failure of inhibitory synaptic transmission could underlie a major mechanism of seizure generation in *SCN8A* epileptic encephalopathy.

### Elevated sodium currents in *Scn8a* mutant PV interneurons

We observed an increase in I_NaP_ in both *Scn8a*^D/+^ and *Scn8a*^W/+^-PV interneurons with an increase in I_NaR_ in *Scn8a*^W/+^-PV interneurons. However, we observed no difference in the transient sodium current in *Scn8a*^D/+^ and *Scn8a*^W/+^-PV interneurons, although it is possible that excised somatic patches may not have recapitulated the high levels of Na_v_1.6 in the axon. Previous studies suggest that *Scn8a* may have a much larger role in resurgent current than transient current.^45^ Increases in I_NaP_ have been implicated in various epilepsies,^42,46,48,71^ and prior computational modeling suggests that heightened I_NaP_ underlies the premature depolarization block phenotype in inhibitory interneurons.^11^ I_NaP_ also functions as an amplifier of synaptic currents,^72,73^ although we did not observe differences in amplitude of uIPSCs in recordings of synaptically-connected pairs. Because I_NaP_ is a consistent, non-inactivating component of the sodium current,^48^ we hypothesize that elevations in I_NaP_ contribute to premature failure of PV interneurons and subsequent entry into depolarization block. Additionally, I_NaR_ currents are crucial in facilitating repetitive, high-frequency firing as they impact fast inactivation through an open channel block,^44,74^ and Na_v_1.6 is a crucial contributor to I_NaR_.^45^ We only observed a significant increase in I_NaR_ in *Scn8a*^W/+^-PV interneurons and not in *Scn8a*^D/+^ PV interneurons, possibly due to mutation-specific effects: this has been observed previously in patient-derived neurons.^75^ Increases in I_NaR_ likely provide excessive depolarizing current resulting in an increase in firing frequencies, which may be responsible for differences observed between *Scn8a*^D/+^ and *Scn8a*^W/+^-PV interneuron firing, as *Scn8a*^D/+^ PV interneurons enter depolarization block at lower current injections.

### Implications for *SCN8A* epileptic encephalopathy

Patients with *SCN8A* mutations are typically treated with sodium channel blockers and many are refractory to treatment, highlighting the need to further understand the basic mechanisms surrounding the *SCN8A* epileptic encephalopathy phenotype. Hyperexcitability of excitatory neurons has often been suggested as the underlying cause behind seizures in *SCN8A* EE, and, contradictory to our results here, a previous study suggests limited involvement of inhibitory interneurons due to the lack of seizures when the R1872W *SCN8A* mutation is expressed in all inhibitory interneurons.^28^ In this study, we provide compelling support for a major involvement of PV inhibitory interneurons in the onset of spontaneous seizures and seizure-induced death in *SCN8A* EE. Gene therapies are in development for both *SCN8A* EE and Dravet Syndrome, and downregulation of *Scn8a* has been shown to reduce seizures in both disorders.^76–78^ Our previous studies have shown that antisense oligonucleotide-mediated rescue of PV interneuron firing reduces seizures and prevents SUDEP in a model of Dravet Syndrome^78^; a similar phenotype may be observed in *SCN8A* EE where rescue of depolarization block prevents seizures and SUDEP. In a similar manner to Dravet Syndrome, specific targeting of inhibitory interneurons in *SCN8A* EE may be a novel therapeutic strategy.

In conclusion, we show here that PV interneurons play a significant role in *SCN8A* epileptic encephalopathy. Elevations in I_NaP_ current likely render PV interneurons more susceptible to action potential failure, and subsequent depolarization block leads to a decrease in network inhibition. PV interneurons also exhibit impaired synaptic transmission, and altogether, we observe that a gain-of-function *SCN8A* mutation exclusively in PV interneurons conveys susceptibility to spontaneous seizures and SUDEP. In the field of *SCN8A* EE, prior research has focused primarily on the impact of GOF *SCN8A* mutations on excitatory neurons.^10,28,79^ These results, along with our previous work proposing that SST interneurons contribute to seizures,^11^ shift the paradigm of the *SCN8A* epileptic encephalopathy field from primarily considering excitatory neuron hyperexcitability as the main driver of the seizure phenotype and calls for future studies to further explore the importance of inhibitory neuron activity in *SCN8A* epileptic encephalopathy.

## Acknowledgements

We thank Miriam Meisler, Ph.D at the University of Michigan for the gift of the *Scn8a*^D/+^ and *Scn8a*^W/+^ mice.

## Data availability

Data will be made available by the corresponding author upon reasonable request.

## Supplementary Data

Supplementary material is available in the online version.

